# The vaginal microbiome drives endometriosis pain

**DOI:** 10.1101/2025.08.21.671604

**Authors:** McKenna L. Pratt, Zulmary Manjarres, Katelyn E. Sadler

## Abstract

Despite being one of two cardinal disease symptoms, endometriosis pain is poorly understood. Using a validated mouse model, we demonstrate that endometriosis-associated vaginal dysbiosis is sufficient to induce pain in the absence of disease pathology. In addition, intravaginal antibiotic treatment and vaginal microbiome transplant from healthy control animals reverses pain in endometriosis mice.

## Main Text

Pain is one of two cardinal symptoms of endometriosis, a disease defined by the presence of endometrial-like tissue outside of the uterus^1^. Historically, endometriosis pain was exclusively attributed to nociceptive signaling in extrauterine lesions. This dogma, however, is challenged by the fact that lesion number and size do not correlate with pain reports and lesion removal does not alleviate pain in a subset of patients^2,3,4^. Given this, additional drivers of endometriosis pain must exist. Previous studies demonstrate that individuals with endometriosis exhibit vaginal dysbiosis^5,6^. Considering the causal role of gut dysbiosis in other chronic pain conditions^7,8^ we investigated the extent to which the vaginal microbiome drives endometriosis pain using mouse models.

Endometriosis was induced in naïve C57BL/6 female mice through intraperitoneal injection of donor uterine tissue^9^. This procedure resulted in the development of extra-uterine lesions throughout the abdominal cavity (**Fig. 1a, 1b**). Relative to sham controls, endometriosis mice developed persistent abdominal mechanical allodynia, a feature also reported in individuals with endometriosis^10,11,12,13^ **Fig. 1c, 1d**; **Extended Data Fig. 1**). Endometriosis mice also developed spontaneous pain as indicated by higher facial grimace scores, more frequent abdominal licking, and reduced rearing behavior (**Fig. 1e-h**). To examine vaginal microbiome contributions to these phenotypes, pain-like behaviors were re-assessed in endometriosis mice following 5 days of intravaginal antibiotic administration (**Fig. 2a**). Elimination of the vaginal microbiome decreased both abdominal mechanical allodynia and ongoing pain in endometriosis mice (**Fig. 2b-d; Extended Data Fig. 1**). These findings suggest that elimination of the dysbiotic vaginal microbiome is sufficient to decrease pain in endometriosis.

**Fig. 1.**
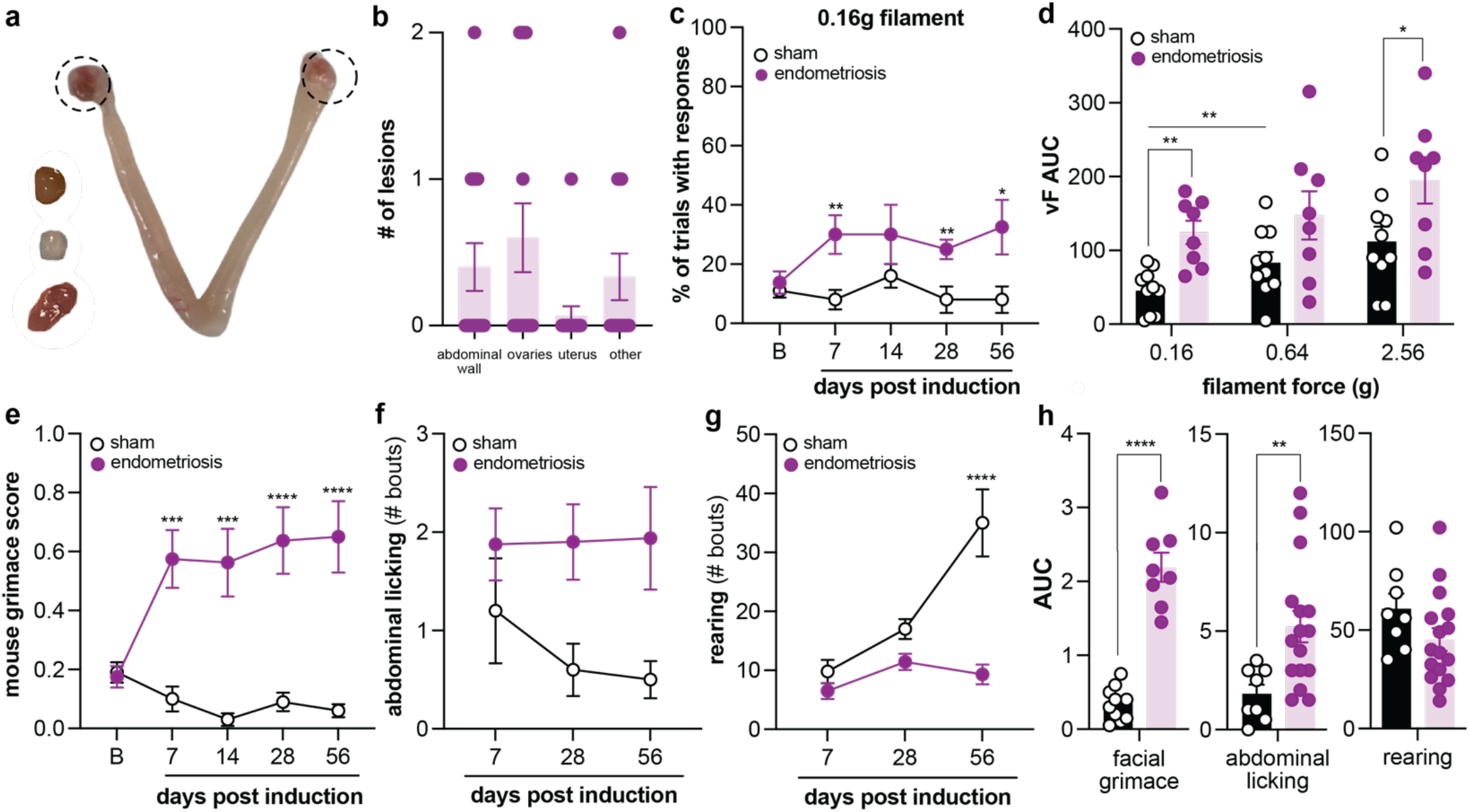
Murine model of endometriosis. **a**, Example endometriotic lesions isolated from intraperitoneal cavity or adhered to ovaries. **b**, Total number of lesions per location from *N*=15 endometriosis mice. **c**, Abdominal mechanical sensitivity of endometriosis and sham mice displayed as a percent of trials with response to the 0.16g von Frey filament from baseline to 56 days post endometriosis/sham induction; *N*=8-10. **d**, Responses to abdominal stimulation with 0.16, 0.64, and 2.56 g von Frey filaments summed from baseline to day 56 post-sham or endometriosis induction. **e**, Facial grimace scores, **f**, bouts of abdominal licking and **g**, bouts of rearing observed in endometriosis and sham mice up to day 56 post endometriosis/sham induction; *N*=8-16. **h**, Ongoing pain behaviors summed from up to day 56 post-sham or endometriosis induction. Percent of trials with response, mouse grimace score, abdominal licking, and rearing comparisons were performed using 2-way repeated measures ANOVA with post hoc Bonferroni correction. von Frey AUC was analyzed using a 2-way mixed ANOVA with post hoc Bonferroni correction. Facial grimace, abdominal licking, and rearing AUC were analyzed using individual two-tailed unpaired t-tests. ^*^*P* ≤ 0.05, ^**^*P* ≤ 0.01, ^***^*P* ≤ 0.001, ^****^*P* ≤ 0.0001.

**Fig. 2.**
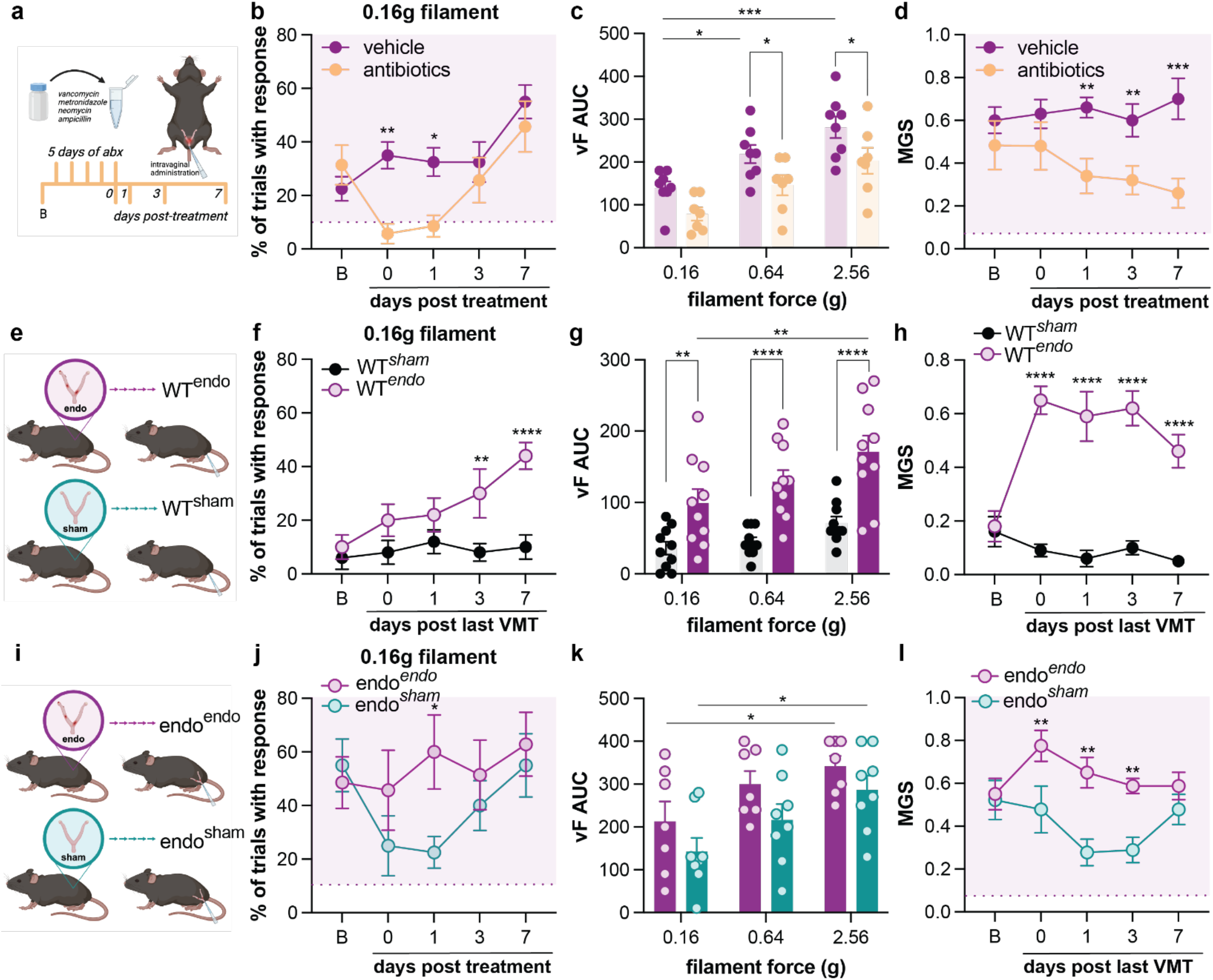
The vaginal microbiome of endometriosis mice is a target for treating pain-like behaviors. **a**, Schematic depicting the timeline and details of antibiotic treatment of endometriosis mice. **b**, Mouse grimace scores of vehicle- and antibiotic-treated endometriosis animals throughout treatment paradigm (*N=*7-8). **c**, Abdominal mechanical allodynia of vehicle- and antibiotic-treated endometriosis mice displayed as a percent of trials with response to a 0.16g von Frey filament from baseline to 7 days post-treatment. **d**, Area under the curve of the graph in (c) along with area under the curve of von Frey filaments 0.64g and 2.56g. **e**, Schematic of VMT from endometriosis to wildtype mice. **f**, Mouse grimace scores of sham and endometriosis VMT wildtype recipients from baseline to 7 days post-treatment (*N=*10). **g**, Abdominal mechanical allodynia of sham and endometriosis VMT recipients displayed as a percent of trials with response to a 0.16g von Frey filament throughout the treatment paradigm. **h**, Area under the curve of the graph in (g) along with area under the curve of von Frey filaments 0.64g and 2.56g. **i**, Schematic of sham VMT to endometriosis mice. **j**, Mouse grimace scores of sham VMT endometriosis mice from baseline to 7 days post-VMT (*N=*7-8). **k**, Abdominal mechanical allodynia of sham VMT endometriosis mice displayed as a percent of trials with response to a 0.16g von Frey filament throughout the treatment paradigm. **l**, Area under the curve of the graph in (k) along with area under the curve of von Frey filaments 0.64g and 2.56g. Percent of trials with response and mouse grimace scores were compared using 2-way repeated measures ANOVA with post hoc Bonferroni correction. Area under the curve for von Frey stimulation response was compared using a 2-way mixed effects ANOVA with post hoc Bonferroni correction. ^*^*P* ≤ 0.05, ^**^*P* ≤ 0.01, ^***^*P* ≤ 0.001, ^****^*P* ≤ 0.0001. Dotted lines indicate the mean percent trials with response and mouse grimace score for sham animals.

Next, we performed a series of vaginal microbiome transplant (VMT) experiments to assess if endometriosis dysbiosis induces pain in the absence of disease pathology (**Fig. 2e**). Five consecutive days of VMT were performed from endometriosis or sham donor mice to naïve C57BL/6 recipients. Resulting changes in pain-like behaviors were recorded for up to one week following VMT completion. Endometriosis VMT recipients developed abdominal mechanical allodynia and ongoing pain that mirrored phenotypes observed in VMT donors (**Fig. 2f-h; Extended Data Fig. 1**). These results support the idea that endometriosis-associated dysbiosis is sufficient to induce persistent pain, even in the absence of ectopic lesions. In a reciprocal fashion, we finally assessed whether introduction of a healthy vaginal microbiome reverses endometriosis pain. For these studies, endometriosis mice received five consecutive days of VMT from sham controls or a separate cohort of mice with endometriosis (**Fig. 2i**). Sham VMT resulted in a transient alleviation of abdominal mechanical allodynia and ongoing pain in endometriosis mice (**Fig. 2j-l; Extended Data Fig. 1**). Thus, introduction of probiotic vaginal species should be explored as a novel means of treating endometriosis pain.

To our knowledge, this study is the first to provide a direct link between the vaginal microbiome and endometriosis pain. Our work demonstrates that both intravaginal antibiotic treatment and VMT from healthy control mice temporarily alleviates pain in mice with endometriosis. The transient nature of this pain relief suggests that disease-specific selective pressures are not modified by either treatment. Regardless, these data provide a foundation on which analgesics based on microbial compounds – a new class of therapies for endometriosis treatment – can be developed. This is particularly notable given that current first-line treatments for endometriosis pain including hormonal contraceptives and non-steroidal anti-inflammatory drugs fail to provide sufficient pain relief for many patients^14,15^. Limitations of the current study include species differences in the vaginal microbiome; the human vaginal microbiome is almost exclusively colonized by *Lactobacilli* whereas the mouse vaginal microbiome is more diverse^16,17,18,19,20,21^. As such, future translational investigations should examine convergent microbial metabolic pathways in the vagina that are disrupted in both mouse models and patients with endometriosis.

## Acknowledgements

This work was funded by grants from the National Institutes of Health (R00HL155791 to KES) and the Rita Allen Foundation (Award in Pain to KES). The authors thank Nurfatihah Zulkifly for assistance with spontaneous behavior video scoring.

## Author contributions

MLP, ZM, and KES conceptualized study and developed methodology. MLP and ZM performed the investigation. MLP and KES are responsible for data visualization. MLP wrote the original manuscript draf; ZM and KES editied draft. KES acquired funds and supervised the project.

## Methods

### Animals

Female C57BL/6 (Jackson Laboratory) mice were bred in house and maintained on a 12-hour light/12-hour dark schedule with *ad libitum* access to rodent chow and water. Animals were 7-to 8-weeks-old at induction of endometriosis; similar-aged mice were used as uterine tissue donors and endometriosis VMT recipients. All protocols were in accordance with the National Institutes of Health guidelines and were approved by the Institutional Animal Care and Use Committee at the University of Texas at Dallas (Richardson, TX; 2022-0088).

### Non-surgical mouse model of endometriosis

Endometriosis was induced using procedures identical to those described by Fattori and colleagues9. Briefly, uterine horn donors received a single injection of 3 μg/100 μL ≤-estradiol 3-benzoate (E2) 7 days prior to tissue collection. E2 was prepared by dissolving 1 mg of E2 (Sigma Aldrich) in 1 mL of 100% molecular grade ethanol (Sigma Aldrich) to a final stock concentration of 1 mg/mL. To create a working solution, 3.3 μL of stock solution was added to 106.7 μL of sesame oil (Sigma Aldrich) such that the final concentration was 3 μg/100 μL. Using a 1cc syringe (Medline), 100 μL of the working solution was injected subcutaneously in each donor mouse. 7 days post-injection, donor mice were sacrificed, and their uterine horns were removed. Uterine horns were cut longitudinally with a pair of dissecting scissors and then minced using scissors and a blade until fragments were < 1 mm in length. Further homogenization of tissue fragments was achieved using a dounce tissue homogenizer (Grainger). Homogenized tissue from all donors was combined into one solution prior to transplant to decrease potential inter-individual variability effects; tissue from 1 donor animal was needed for every 2 recipients. Recipient mice were maintained on inhalable isoflurane (1.8-2.2%; Covetrus) for the duration of induction. Using a 23G needle (PrecisionGlide™), 500 μL of uterine horn solution was injected intraperitoneally into each recipient mouse. Sham mice were given an intraperitoneal injection of 500 μL of Hank’s balanced salt solution. Mice were immediately returned to their home cage following induction.

### Vaginal lavage

Vaginal lavage was obtained as previously described22. Briefly, mice were scruffed then, using a p200 pipette, 50 μL of sterile saline (Medline) was flushed in and out of the vagina 5 times. Samples were collected in sterile tubes and either frozen at -80°C or placed on ice for immediate use.

### Behavioral tests

Prior to the start of all behavioral tests, mice were habituated to the testing apparatus for 2 hours and experimenter presence for 30 minutes. The experimenter was blinded to injury and drug treatment where applicable. Mice were randomly assigned to all treatment groups.

### Spontaneous behaviors

Spontaneous behavior was hand scored from 10-minute bottom-up videos. Videos were taken prior to other behavior testing at each timepoint. Scored behaviors included rearing and abdominal or non-abdominal licking/grooming behaviors. Rearing behavior was classified as the lifting of both front paws from the ground and standing in an upright position on one or both hindlimbs. Supported rearing (against the walls of the behavior chamber) and unsupported rearing were considered equal behaviors. For spontaneous abdominal pain measurements, licking of the abdomen was considered and analyzed as previously reported9. Non-abdominal licking was also analyzed. For each spontaneous behavior measured, the number of bouts per activity was reported.

### Mouse grimace score

Facial grimace was scored manually by a trained experimenter as previously described23. This was measured following the completion of bottom-up video recording and prior to further behavior testing. Mice were given a score of 0-2 for 5 different facial features: orbital tightening, nose bulge, cheek bulge, ear position, and whisker change. Each animal was scored twice with approximately 5 min between trials. The scores for all features were averaged to generate the overall mouse grimace score.

### von Frey mechanical allodynia testing

Abdominal sensitivity to punctate mechanical stimulation was assessed via calibrated von Frey filaments (0.16g; 0.64g; 2.56g)^24^. These filaments were selected based on preliminary ‘up/down’ abdominal von Frey testing completed in sham animals; the average 50% withdrawal threshold in these mice was 0.64g. Thus, we included the 0.64g filament and 2 filaments ‘up’ and ‘down’ from that average: 0.16g and 2.56g. The decision to include 3 filaments was made to better capture a mechanical allodynia phenotype and/or a change in that phenotype. Animals’ abdomens were stimulated 5 times with each filament with at least 1 minute between stimulations. Response frequency was recorded for each filament. Care was taken to avoid stimulating the same spot on the abdomen twice. A jump, paw flinch, or kicking of hindpaws was counted as a withdrawal response.

### Vaginal treatments

Volumes for all vaginal treatments (25 μL) was determined through pilot studies wherein various amounts of crystal violet (1%) was administered into the vaginal cavity of mice. N=3 mice were used for each volume tested (10 μL; 25 μL; 50 μL). Mice were placed supine and maintained on isoflurane (1.8-2.2%) during the treatment. Following administration, 5 minutes were allowed for the crystal violet solution to sit in the vaginal cavity prior to removing the mouse from isoflurane and immediately euthanizing. This 5-minute period was provided for all treatments to avoid immediate leakage of the solution out of the vagina. In crystal violet pilot studies, the vaginal tissue was dissected from each mouse and cut open longitudinally. Images were taken and analyzed using ImageJ for percent coverage (Extended Figure 2).

### Antibiotic treatment

Antibiotic cocktail was made fresh every other day of treatment. This cocktail is commonly used to create a pseudo germ-free intestine in rodents and contains 4 antibiotics: vancomycin (0.5 g/L), ampicillin (1 g/L), neomycin (1 g/L), and metronidazole (1 g/L)25. To make the solution, antibiotics were resuspended in sterile saline at the appropriate concentrations. Mice were placed supine and maintained on inhalable isoflurane (1.8-2.2%) during treatment. 25 μL of the antibiotic cocktail was intravaginally administered using sterile 200 μL pipet tips. Vehicle animals received 25 μL of sterile saline. Following administration, mice remained under isoflurane for 5 minutes. Animals were immediately returned to their home cage following the 5-minute absorption period and closely observed until normal behavior resumed. Treatment was repeated daily for 5 consecutive days.

### Vaginal microbiome transplant

Vaginal lavage contents were obtained from donors using the lavage method described above. To avoid potential animal and/or cage-specific effects, vaginal lavage donors were rotated in groups such that no donor was used on two consecutive days. This rotation scheme was also used to reduce potential lavage-related disruptions in donor vaginal microbiomes. 50 μL of lavage contents were collected from each donor mouse and combined to create a mixed donor solution. This solution was stored on ice throughout lavage collection and vaginal microbiome transplant (VMT). VMT was performed immediately following collection. Recipient mice were placed supine and maintained on inhalable isoflurane (1.8-2.2%) during treatment. 25 μL of the pooled vaginal lavage contents were intravaginally administered using sterile 200 μL pipet tips. Following administration, mice remained under isoflurane for 5 minutes before being returned to their home cage. Mice were observed closely until normal behavior resumed. VMTs were performed daily for 5 consecutive days.

### Extra-uterine lesion detection

At the end of each animal’s treatment paradigm, mice were humanely sacrificed using isoflurane overdose followed immediately by cervical dislocation. The abdominal region of mice with endometriosis mice was carefully examined for ectopic lesions. The entire length of the abdominal wall, uterine horns, ovaries, and all other tissue located within the peritoneal cavity were searched for endometrial adhesions. Location and number of lesions in each endometriosis mouse was recorded to confirm success of the disease model.

### Statistical analysis

Individual data points were presented wherever possible in addition to group means ± SEM. Data were analyzed using GraphPad Prism 10; results were considered statistically significant when *P* < 0.05.

---

**Extended Data Fig. 1.**
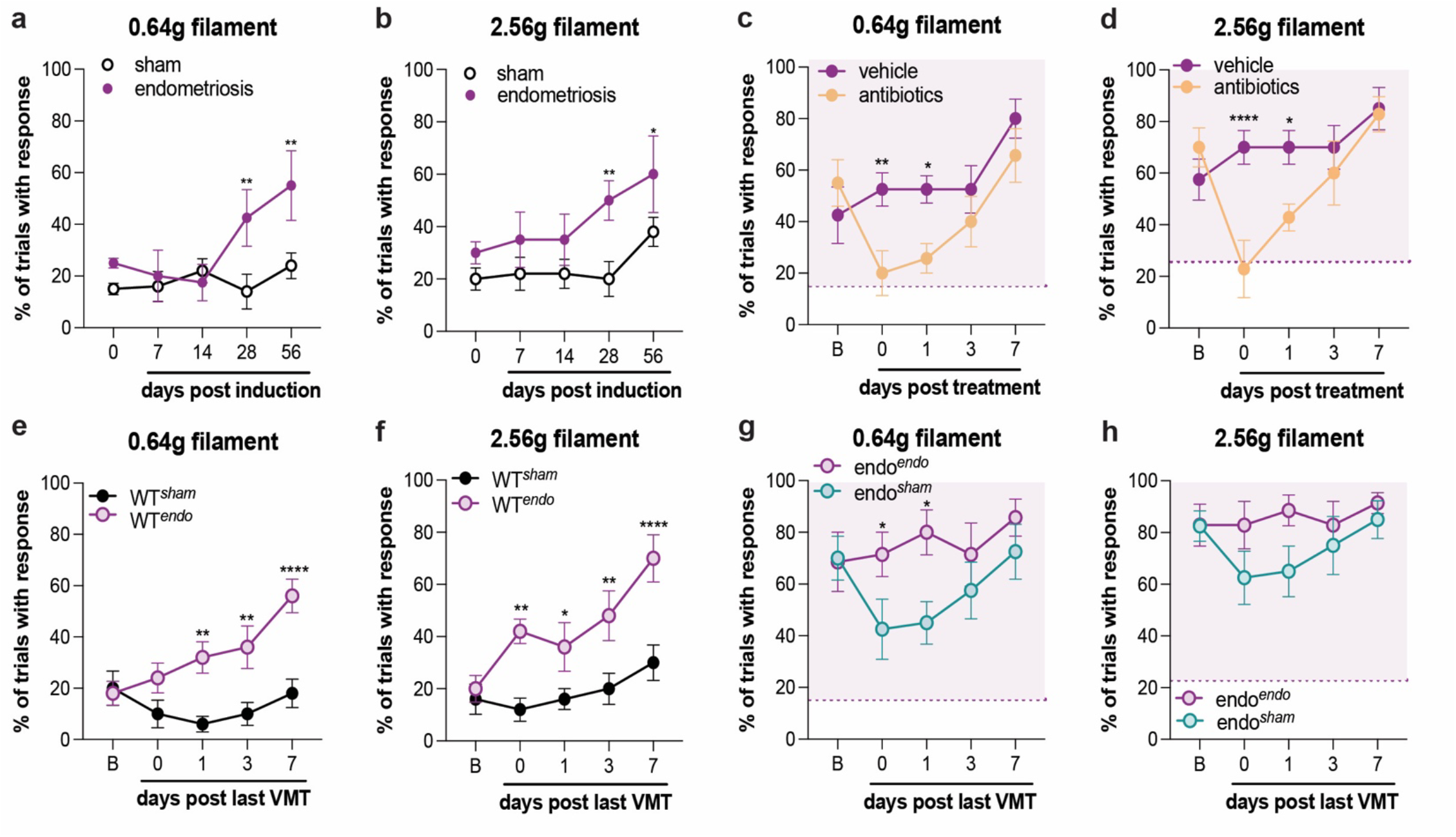
Additional abdominal mechanical hypersensitivity tests. Percent of trials with response to the 0.64g and 2.56g von Frey filament for sham versus endometriosis mice (**a, b**; *N=*8-10), vehicle versus antibiotic-treated endometriosis mice (**c, d**; *N=*7-8), sham versus endometriosis VMT-treated wildtype (WT) recipients (**e, f**; *N=*10), and sham versus endometriosis VMT-treated endometriosis mice (**g, h**; *N=*7-8). All comparisons were performed using a 2-way repeated measures ANOVA with post hoc Bonferroni correction. ^*^*P* ≤ 0.05, ^**^*P* ≤ 0.01, ^***^*P* ≤ 0.001, ^****^*P* ≤ 0.0001. Dotted lines indicate the mean percent trials with response for sham animals.

**Extended Data Fig. 2.**
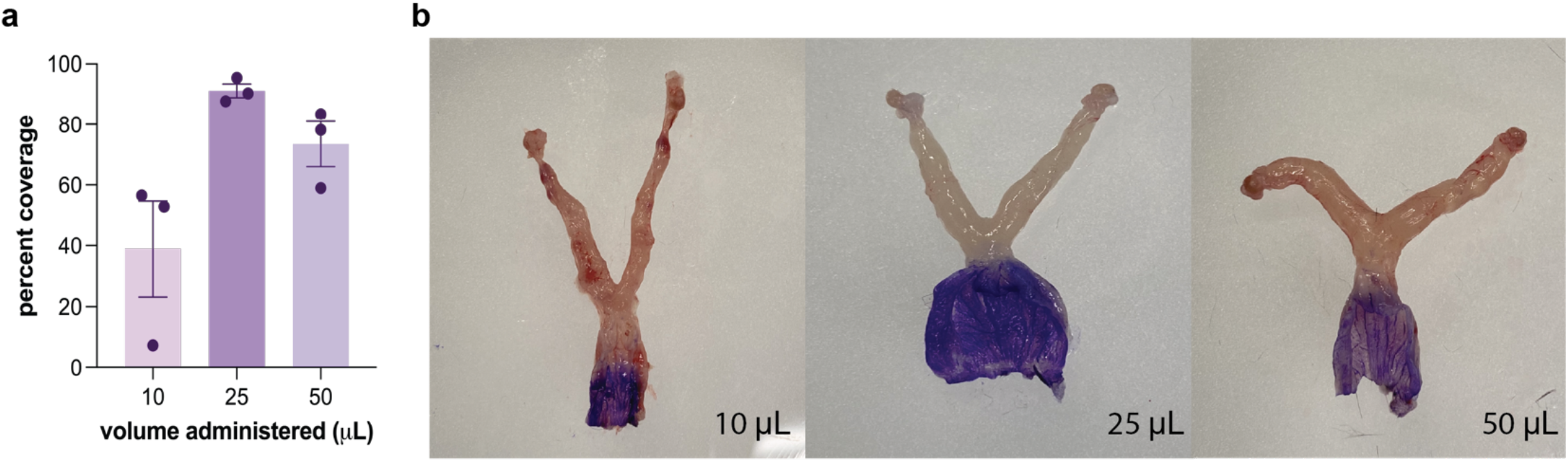
Pilot study for volume of vaginal treatments. **a**, Plot showing percent coverage by 3 different volumes of crystal violet (1%) administered to the vagina of mice (*N=*3 per volume group). **b**, Representative images of 3 reproductive tracts following intravaginal administration of 10, 25, or 50 μL of crystal violet (1%). Tissue was cut longitudinally to expose the vaginal cavity.

