## Extended Figures for "The vaginal microbiome drives endometriosis pain"

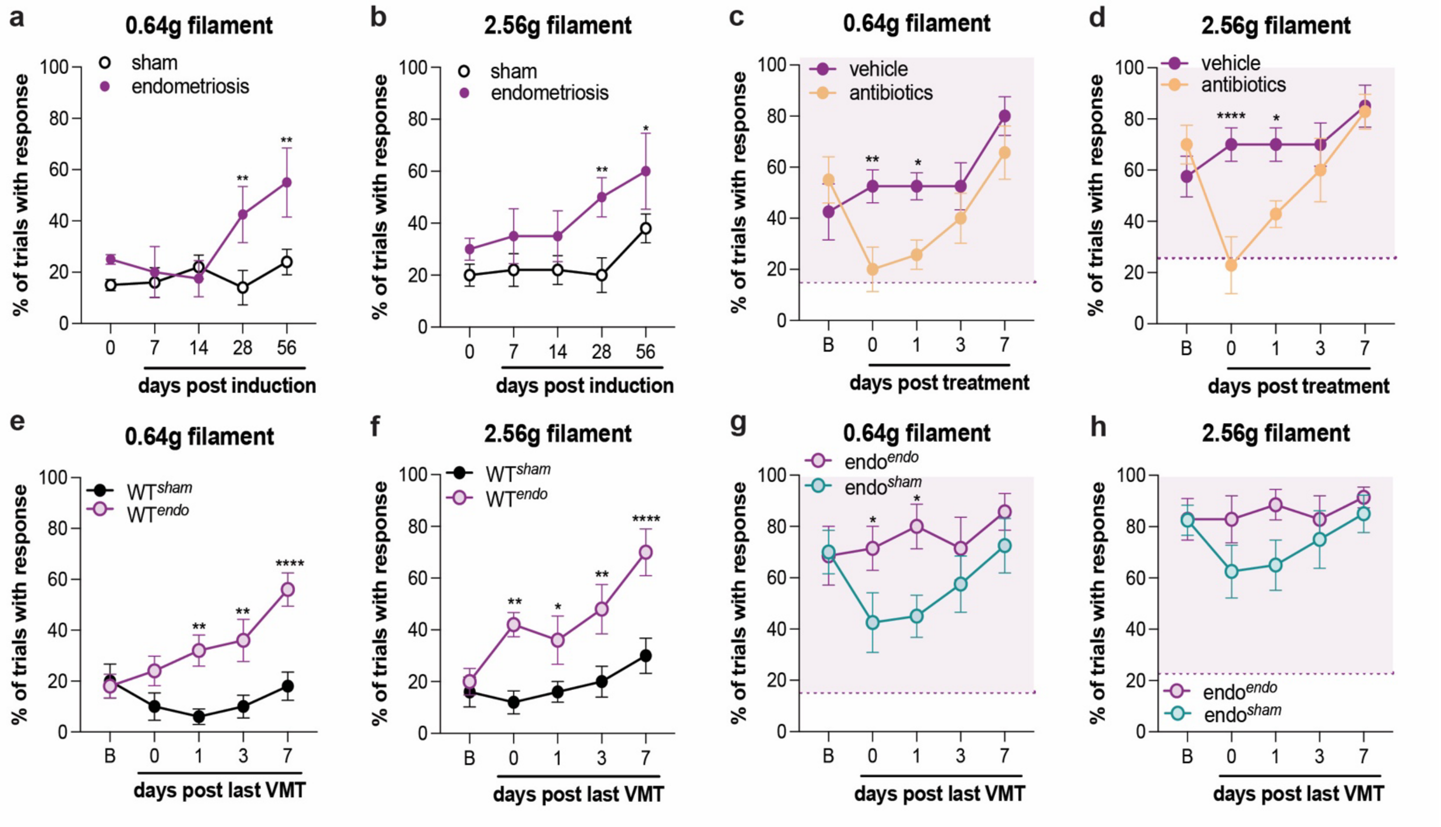


**Extended Data Fig. 1 | Additional abdominal mechanical hypersensitivity tests.** Percent of trials with response to the 0.64g and 2.56g von Frey filament for sham versus endometriosis mice (**a, b**; $N$=8-10), vehicle versus antibiotic-treated endometriosis mice (**c, d**; $N$=7-8), sham versus endometriosis VMT-treated wildtype (WT) recipients (**e, f**; $N$=10), and sham versus endometriosis VMT-treated endometriosis mice (**g, h**; $N$=7-8). All comparisons were performed using a 2-way repeated measures ANOVA with post hoc Bonferroni correction. $^{*}P \leq 0.05$, $^{**}P \leq 0.01$, $^{***}P \leq 0.001$, $^{****}P \leq 0.0001$. Dotted lines indicate the mean percent trials with response for sham animals.

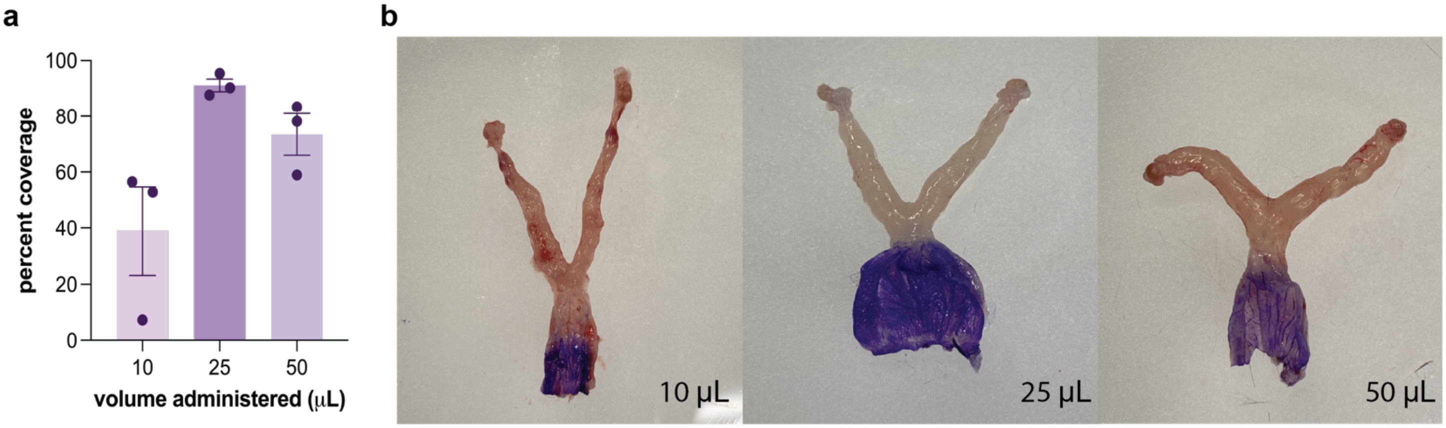


**Extended Data Fig. 2 | Pilot study for volume of vaginal treatments. a,** Plot showing percent coverage by 3 different volumes of crystal violet (1%) administered to the vagina of mice ($N$=3 per volume group). **b,** Representative images of 3 reproductive tracts following intravaginal administration of 10, 25, or 50 μL of crystal violet (1%). Tissue was cut longitudinally to expose the vaginal cavity.
